# Geographical Survey of the Mycobiome and Microbiome of Southern California Glassy-winged Sharpshooters

**DOI:** 10.1101/2023.04.27.538478

**Authors:** Cassandra L. Ettinger, Jessica Wu-Woods, Tania L. Kurbessoian, Dylan J. Brown, Inaiara de Souza Pacheco, Beatriz G. Vindiola, Linda L. Walling, Peter W. Atkinson, Frank J. Byrne, Richard Redak, Jason E. Stajich

## Abstract

The glassy-winged sharpshooter, *Homalodisca vitripennis* Germar, is an invasive xylem-feeding leafhopper with a devastating economic impact on California agriculture through transmission of the plant pathogen, *Xylella fastidiosa*. While studies have focused on *X. fastidiosa* or known symbionts of *H. vitripennis*, little work has been done at the scale of the microbiome (the bacterial community) or mycobiome (the fungal community). Here we characterize the mycobiome and the microbiome of *H. vitripennis* across Southern California and explore correlations with captivity and host insecticide-resistance status. Using high-throughput sequencing of the ribosomal internal transcribed spacer (ITS1) region and the 16S rRNA gene to profile the mycobiome and microbiome, respectively, we found that while the *H. vitripennis* mycobiome significantly varied across Southern California, the microbiome did not. We also observed a significant difference in both the mycobiome and microbiome between captive and wild *H. vitripennis*. Finally, we found that the mycobiome, but not the microbiome, was correlated with insecticide-resistance status in wild *H. vitripennis*. This study serves as a foundational look at the *H. vitripennis* mycobiome and microbiome across Southern California. Future work should explore the putative link between microbes and insecticide-resistance status and investigate whether microbial communities should be considered in *H. vitripennis* management practices.

## Introduction

*Homalodisca vitripennis* Germar, the glassy-winged sharpshooter (GWSS), is an invasive xylem-feeding leafhopper with a broad host range in California spanning over 340 plant species (https://www.cdfa.ca.gov/pdcp/Documents/HostListCommon.pdf). GWSS is the primary vector in California of the bacterial pathogen, *Xylella fastidiosa* Wells, the causal agent of several important diseases in economically important agricultural plants including grapes, peaches, citrus, and almonds [1]. Like many xylem-feeding insects, GWSS relies on two obligate bacterial symbionts, *Candidatus* Sulcia muelleri and *Ca*. Baumannia cicadellinicola, for biosynthesis of essential amino acids, which are limited in its xylem-based diet [2–8]. Additionally, the facultative symbiont, *Wolbachia* sp., has been reported as abundant in GWSS [2, 8–13]. Despite detailed investigations into the obligate bacterial symbionts of GWSS and its association with *X. fastidiosa*, comparatively little is known about the overall microbiome of GWSS, and little has been reported about the composition of the mycobiome.

Native to the southeastern USA and northeastern Mexico, GWSS was introduced to California in the 1990s [14–16]. Since its introduction, area-wide treatments of insecticides, particularly the systemic neonicotinoid insecticides, imidacloprid and acetamiprid, have been used to control these invasive insects with some success [17, 18]. However, starting in 2012 the effectiveness of population control by these insecticides appeared substantially weakened [19, 20], with documented instances of neonicotinoid applications leading to high levels of insecticide-resistance in some Southern California populations resulting in GWSS population resurgence [21, 22]. insecticide-resistance usually involves multiple coexisting mechanisms spanning behavioral (e.g. avoidance) and physiological processes (e.g. cuticle modifications, detoxification) [23–25], and studies have proposed a novel role for symbionts and other associated microbes in detoxification of insecticides for their associated hosts [26–31]. Recent studies on Southern California GWSS populations have identified both trade-offs in host reproductive fitness associated with resistance [32] as well as host genes that may play a role in conferring resistance [33]. However, the possible role of the microbiome and mycobiome in the resistance of these GWSS populations has yet to be fully explored.

Bacteria can have critical functional roles that affect host insect fitness, ranging from pathogenicity to positive benefits such as enhancing nutrient acquisition (e.g. obligate symbionts), protection from pathogens or other stressors via detoxification of phytotoxins and insecticides [34, 35]. Previous GWSS microbiome studies have reported bacterial communities dominated by obligate symbionts, followed by members of the genera, *Wolbachia*, *Xylella*, *Cardiobacterium, Pectobacterium, Serratia*, *Pseudomonas*, *Pantoea*, *Ralstonia*, *Bacillus*, *Pedobacter*, *Methylobacterium*, and *Curtobacterium* [11, 13, 36–38]. While the functional role of many of these taxa has yet to be elucidated, these foundational studies suggest major factors affecting the composition of the GWSS microbiome may include geography, host plant and insect developmental stage [36, 37].

In contrast to bacteria, fungi are an underappreciated part of insect-associated microbial communities [35] and there have been no studies profiling the GWSS mycobiome using culture-independent approaches. Instead, previous studies have focused on identification of entomopathogenic fungi that can infect GWSS for use in population management through biocontrol including *Hirsutella homalodiscae*, *Pseudogibellula formicarum*, *Metarhizium anisopliae*, *Sporothrix* sp., *Beauveria bassiana*, and *Isaria poprawskii* [39–44]. Fungi can inhabit multiple ecological niches, and not all insect-associated fungi are pathogens [45, 46]. For example, yeast-like symbionts (YLS) have been previously identified in the fat body of other Hemipteran insects including cicadas, scales and planthoppers [47–51]. These YLS can even provide nutritional benefits to planthoppers given their nutritionally limited diets [52]. While YLS have not been reported yet in GWSS, it is possible that fungal community members are capable of forming similarly important and complex roles in association with GWSS.

Given what little is known about the factors shaping the microbial communities of this invasive pest insect, we characterized the taxonomic composition of the mycobiome and the microbiome of GWSS in Southern California using high-throughput amplicon sequencing. We addressed three ecological questions related to these microbial communities: (i) do these communities vary across geographic regions, (ii) does captivity correlate with a shift in these communities, and (iii) can we detect a microbial signal correlated with host insecticide-resistance status in wild GWSS that might be useful for understanding and identifying resistance of the host insect population.

## Methods

### Sample collection

Wild-caught GWSS were collected between 2017-2022 during insecticide-resistance monitoring work (i.e., [21]) from nine populations spanning six geographic regions across Southern California (<u>Figure S1</u>). Captive lines representing GWSS collected from three of these populations were maintained and individuals from these lines were sacrificed in 2022 for use here. In total, 87 GWSS were sampled, representing 63 opportunistically caught wild GWSS and 24 captive GWSS (Table 1). In addition to captive GWSS from the three long-term captive lines, we also sampled members of a newly established line of first generation (G1) offspring from a wild-caught individual from Riverside. GWSS were sexed during collection or prior to processing (female = 49, male = 38). The imidacloprid-resistance status is known for 53 of these GWSS from assays of these individuals themselves or other individuals from the same collection date or captive line. Imidacloprid status assays were performed as described in Ettinger et al [33]. Insects were stored in RNA*later* (ThermoFisher Scientific, Waltham, MA, USA) or 200 proof ethanol and kept at −20°C prior to processing.

**Table 1.**
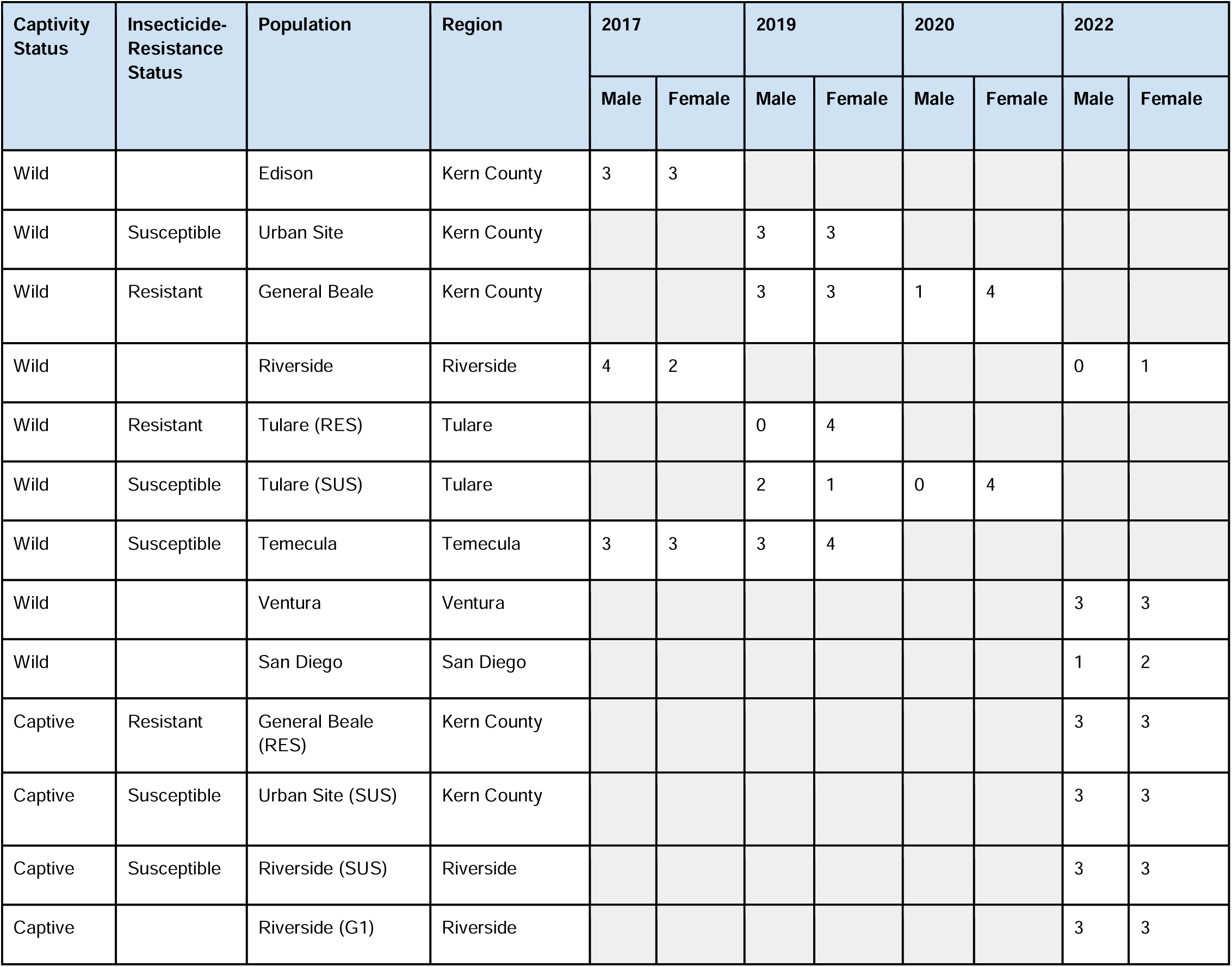
Summary table describing the number of GWSS sequenced in this work. This table reports on the number of male and female GWSS collected in each year and their captivity status, host imidacloprid resistance status, the collection site (population), and collection region.

| Captivity Status | Insecticide-Resistance Status | Population | Region | 2017 |  | 2019 |  | 2020 |  | 2022 |  |
| --- | --- | --- | --- | --- | --- | --- | --- | --- | --- | --- | --- |
|  |  |  |  | Male | Female | Male | Female | Male | Female | Male | Female |
| Wild |  | Edison | Kern County | 3 | 3 |  |  |  |  |  |  |
| Wild | Susceptible | Urban Site | Kern County |  |  | 3 | 3 |  |  |  |  |
| Wild | Resistant | General Beale | Kern County |  |  | 3 | 3 | 1 | 4 |  |  |
| Wild |  | Riverside | Riverside | 4 | 2 |  |  |  |  | 0 | 1 |
| Wild | Resistant | Tulare (RES) | Tulare |  |  | 0 | 4 |  |  |  |  |
| Wild | Susceptible | Tulare (SUS) | Tulare |  |  | 2 | 1 | 0 | 4 |  |  |
| Wild | Susceptible | Temecula | Temecula | 3 | 3 | 3 | 4 |  |  |  |  |
| Wild |  | Ventura | Ventura |  |  |  |  |  |  | 3 | 3 |
| Wild |  | San Diego | San Diego |  |  |  |  |  |  | 1 | 2 |
| Captive | Resistant | General Beale (RES) | Kern County |  |  |  |  |  |  | 3 | 3 |
| Captive | Susceptible | Urban Site (SUS) | Kern County |  |  |  |  |  |  | 3 | 3 |
| Captive | Susceptible | Riverside (SUS) | Riverside |  |  |  |  |  |  | 3 | 3 |
| Captive |  | Riverside (G1) | Riverside |  |  |  |  |  |  | 3 | 3 |

### Molecular methods and sequence generation

DNA was extracted from GWSS and control samples (n_GWSS_ = 79, n_control_ = 6) using a DNeasy PowerSoil DNA Isolation kit (Qiagen, Germany) with minor changes to the manufacturer’s protocol as follows. To improve fungal lysis, samples were heated at 70°C for 10 min after adding C1 solution. Instead of bead beating, samples were vortexed for 10 minutes following manufacturer instructions. Finally, samples were eluted in only 50 µL of C6 solution. Insects were removed from 1.5-mL tubes using flame sterilized tweezers, sexed and entire insect bodies were placed directly into DNeasy PowerSoil bead tubes prior to DNA extraction. No surface sterilization was performed as it has been previously reported to not affect insect microbiome community structure [53]. Samples were placed into four randomized blocks prior to DNA extraction using a random number generator. DNA extraction was also performed on three no-sample added (negative) and three ZymoBIOMICS Microbial Community Standard (positive) controls (Zymo Research, Irvine, CA, USA). DNA from an additional eight GWSS had been previously extracted using the Blood and Tissue kit (Qiagen) according to the manufacturer instructions.

Using a random number generator, DNA extracts were randomly assigned places in a 96-well plate. Three wells for PCR negative controls (no DNA added) were included in the 96-well plate design. The ribosomal internal transcribed spacer 1 (ITS1) region was amplified using the fungus specific Earth Microbiome Project (EMP) IT1F and ITS2 primer set [54, 55] and the 16S ribosomal RNA (rRNA) gene was amplified using the EMP 515F (Parada) and 806R (Apprill) primer set [56, 57]. PCRs were performed using Platinum Hot Start PCR Master Mix (2x) (ThermoFisher Scientific). For the ITS1 amplicon, duplicate PCRs for each sample were performed in 96-well plate format using the EMP protocol conditions: 94°C for 1 min, 35 cycles at 94°C for 30 s, 52°C for 30 s, 68°C for 30 s, and a final extension at 68°C for 10 min. For the 16S rRNA amplicon, duplicate PCRs for each sample were performed in 96-well plate format using modified EMP protocol conditions: 94°C for 3 min, 35 cycles at 94°C for 45 s, 78°C for 10 s, 50°C for 60 s, 72°C for 90 s, and a final extension at 72°C for 10 min. PCR conditions of the 16S rRNA gene were modified from the EMP protocol to enable inclusion of a clamping step with mitochondrial PNA (PNABio, Newbury Park, CA, USA) to reduce GWSS mitochondrial amplification. Duplicate PCRs were combined prior to clean up and normalization with a SequalPrep Normalization Plate kit (ThermoFisher Scientific) following manufacturer instructions using 25 µL of PCR product per sample. After clean up and normalization, 5 µL from each well was pooled to make the final library for sequencing. Libraries were sequenced at the University of California, Riverside Genomics Core Facility on an Illumina MiSeq (Illumina, San Diego, CA, USA). The ITS1 amplicon library was sequenced to produce 250 bp paired-end reads, and the 16S rRNA gene amplicon library was sequenced to produce 300 bp paired-end reads.

### Sequence processing

Primer sequences were removed using cutadapt v. 2.3 [58]. Figaro was run to inform optimal values for max error and truncation parameters prior to running DADA2 in R [59–61]. For the ITS1 amplicon, reads were processed with maxEE=c(3,3) and truncQ=10, but were not truncated further due to ITS1 length variation. For the 16S rRNA gene amplicon, reads were processed with maxEE=c(1,2), truncLen=c(240,111), and truncQ=10. Reads were then denoised and merged using DADA2 to generate count tables of amplicon sequence variants (ASVs). Chimeric sequences were removed using removeBimeraDenovo (1.35% of ASVs for ITS1 region, 1.58% of ASVs for 16S rRNA gene).

After chimera removal, samples had an average read depth of 24,600 (range: 0 - 76,922) for the ITS1 region and an average read depth of 35,947 (range: 855 - 84,253) for the 16S rRNA gene amplicon data. Taxonomy was inferred using the RDP Naive Bayesian Classifier algorithm with the UNITE (v. 9 “fungi”) database for ITS1 region amplicons and the SILVA (v. 138) database for 16S rRNA gene amplicons [62–65].

To identify possible contaminants, we used decontam’s prevalence method with a threshold of 0.5, which will identify ASVs with a higher prevalence in negative controls than in true samples [66]. This threshold identified 36 and 318 possible contaminants in the ITS1 region amplicon and 16S rRNA gene amplicon data, respectively. These contaminant ASVs were removed from the final datasets and negative and positive controls were subsequently removed at this point in the analysis. Further, for the ITS1 region amplicons, all ASVs taxonomically assigned as nonfungal at the domain level were removed. While, for the 16S rRNA gene amplicons, all ASVs assigned as chloroplasts and mitochondria were removed. The resulting count tables contained 1812 ASVs representing 86 GWSS samples for the ITS1 amplicons, and 2971 ASVs representing 87 GWSS samples for the 16S rRNA gene amplicons. One sample (GW007) was dropped when analyzing ITS1 amplicons as it contained 0 reads after processing.

For relative abundance, alpha and beta diversity analyses, count tables were normalized for sequencing depth by subsetting without replacement to 2000 and 6000 sequences per sample for ITS1 and 16S rRNA gene amplicons respectively. This resulted in 11 and 10 samples for the ITS1 and 16S rRNA gene datasets, respectively, being excluded from rarified analyses. Rarefaction depths were chosen after examining rarefaction curves and library sizes to balance maintaining the maximum number of sequences per sample while also minimizing the number of removed samples. When testing hypotheses about biogeography the amplicon datasets were subset to only samples from wild GWSS (*n_ITS_* = 57, *n_16S_* = 56). When testing hypotheses about captivity and sex all GWSS were included (*n_ITS_* = 75, *n_16S_* = 77). Finally, when testing hypotheses about resistance status, only wild GWSS with a known resistance status were used (*n_ITS_* = 30, *n_16S_* = 28).

### Relative abundance

To visualize community composition across captivity and populations, we transformed rarified read counts from all GWSS samples to proportions and collapsed ASVs into taxonomic genera using the tax_glom function in the phyloseq package [67]. For visualization, we agglomerated genera with a mean proportion of less than 5% into a single “Other” category. For visualization, color palettes from the microshades R package were used [68].

### Diversity analyses

Alpha diversity was calculated for the Shannon index using the estimate_richness function in the phyloseq R package. To test for significant differences in alpha diversity across biogeography, captivity, sex and resistance status, we used Kruskal-Wallis tests with 9,999 permutations. *Post hoc* Dunn tests were performed when the Kruskal-Wallis test resulted in a rejected null hypothesis (*P <* 0.05). All *p*-values were corrected for multiple comparisons using the Benjamini-Hochberg method. Alpha and beta diversity analyses were visualized using tidyverse [69], ggplot2 [70], vegan [71] and phyloseq [67] packages in R.

Beta diversity was calculated for the Hellinger distance using the avgdist function in the vegan R package by providing the unrarefied data and the desired rarefaction depth to calculate an average dissimilarity matrix based on 100 random iterations. The ordinate function in the phyloseq R package was then used to generate principle coordinate analysis plots from the resulting dissimilarity matrices. To test for significant differences in beta diversity centroids (i.e., means of each group) across biogeography, captivity, sex and resistance status, we performed permutational multivariate analyses of variance (PERMANOVAs) with 9,999 permutations and by = “margin” using the adonis2 function in the vegan R package. Collection region and year were included as covariates in models testing captivity and resistance, while year was included in models testing biogeography. For significant PERMANOVA results, *post hoc* pairwise PERMANOVAs were performed using the pairwise.adonis function from the pairwiseAdonis package in R [72]. Further, since PERMANOVA results have been reported to be sensitive to dispersion (i.e., variance) differences for unbalanced designs [73], we also used the betadisper and permutest functions from the vegan package in R with 9,999 permutations to test for significant differences in dispersion. *Post hoc* Tukey’s honestly significant difference (HSD) tests were used to assess which pairwise differences drove significant betadisper results. All *p*-values were corrected using the Benjamini-Hochberg method.

### Differential abundance analysis using DESeq2

For differential abundance analyses, unrarefied datasets were used. For questions about captivity, amplicon datasets included all GWSS (*n_ITS_* = 86, *n_16S_*= 87), while for questions about resistance status, only wild GWSS with a known resistance status were used (*n_ITS_*= 34, *n_16S_* = 35). To identify specific ASVs correlated with captivity or resistance status, we used the DESeq2 package on raw read counts to calculate the differential abundance (log_2_ fold change) of ASVs in both data sets [74]. Collection region and year were included as covariates in both models. All *p*-values were corrected using the Benjamini-Hochberg method. ASVs that differed in abundance were visualized using tidyverse [69], ggplot2 [70] and phyloseq [67] packages in R. We also used FUNGuild v.1.1 to assign trophic guilds to ITS1 ASVs whose abundance significantly correlated with captivity or host insecticide-resistance status [75].

## Results

### Taxonomic composition of the myco- and microbiome of GWSS

The mycobiome was largely dominated by Ascomycota, particularly members of the genera *Cladosporium, Alternaria, Fusarium, Acremonium, Ramularia,* and *Neodidymelliopsis* (Figure 1A). Amplicon sequence variants (ASVs) that were unable to be taxonomically classified into genera were also prevalent in the mycobiome. Additionally, one captive line (Kern County [RES]) showed evidence of a possible enrichment of *Golovinomyces*, while three of the captive lines had low relative abundance of *Basidiobolus* sp.

**Figure 1.**
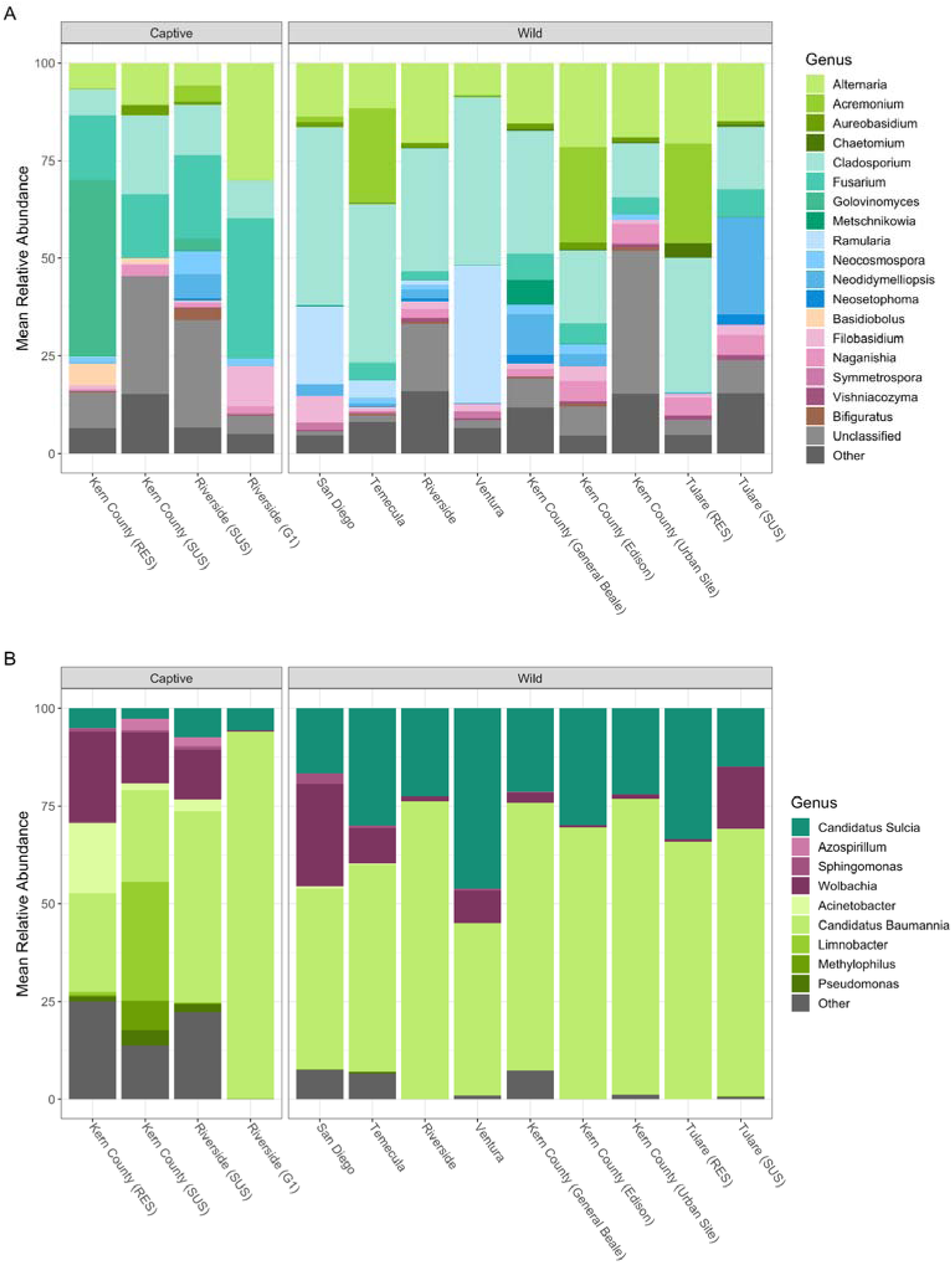
Mean relative abundance of genera associated with captive and wild GWSS populations. Stacked bar charts displaying the mean relative abundance of (A) ITS1 and (B) 16S rRNA gene ASVs for each population (collection site) colored by predicted genera. Genera representing less than 5% mean relative abundance across the dataset are collapsed for visualization purposes into a single group labeled ‘Other’. Number of insects summarized per population are as follows - Kern County (RES): *n_ITS_* = 5, *n_16S_*= 6; Kern County(SUS): *n_ITS_* = 3, *n_16S_* = 5; Riverside (SUS): *n_ITS_* = 5, *n_16S_* = 4; Riverside (G1): *n_ITS_* = 5, *n_16S_*= 6; San Diego: *n_ITS_* = 3, *n_16S_* = 3; Temecula: *n_ITS_* = 12, *n_16S_*= 13; Riverside: *n_ITS_* = 7, *n_16S_* = 4; Ventura: *n_ITS_* = 6, *n_16S_* = 6; Kern County (General Beale): *n_ITS_*= 11, *n_16S_* = 9; Kern County (Edison): *n_ITS_* = 5, *n_16S_* = 6; Kern County (Urban Site): *n_ITS_* = 4, *n_16S_* = 2; Tulare (RES): *n_ITS_* = 3, *n_16S_* = 3; Tulare (SUS): *n_ITS_* = 6, *n_16S_*= 7.

Overall, the microbiome had significantly lower Shannon alpha diversity compared to the mycobiome (K-W test; *p* < 0.01) and was dominated by known obligate symbionts, *Ca.* Baumannia cicadellinicola and *Ca.* Sulcia muelleri, and the facultative symbiont *Wolbachia* sp. (Figure 1B). While facultative, *Wolbachia* sp. were detected in 96.6% of sampled GWSS microbiomes. Despite symbiont dominance, we still observed other bacterial genera at lower relative abundances in the microbiome including *Limnobacter*, *Acinetobacter*, *Methylophilus*, and *Pseudomonas*.

### Only the mycobiome differs across Southern California

Mycobiome beta diversity was significantly different between populations and regions across Southern California as well as over time (Figure S2A; PERMANOVA, *p* < 0.01). Pairwise contrasts found all regional comparisons to be significantly different from each other with two exceptions, San Diego vs. Riverside, and Kern County vs. Tulare (Table S1; *p* > 0.05). Additionally, we found significant differences in dispersion between populations, regions and time (betadisper, *p <* 0.01). These were driven by most regions having higher variance than San Diego and Ventura, possibly due to the smaller sample sizes of those regions (Table S2; Tukey, *p* > 0.05). No significant differences were found in alpha diversity across populations or regions (Figure S3A; Kruskal-Wallis, *p* > 0.05).

While we found differences in the mycobiome that correlated with geographic region, we did not find significant differences in beta diversity across populations, regions or time in the microbiome (Figure S2B; PERMANOVA, *p* > 0.05). There were also no significant differences in dispersion across populations, regions or time (betadisper, *p* > 0.05). Similar to the mycobiome, we did not find any differences in alpha diversity across populations or regions for the microbiome (Figure S3B; Kruskal-Wallis, *p* > 0.05).

### Both the myco- and microbiomes are altered during captivity

We observed significant differences in beta diversity in terms of mean centroids and mean dispersion between captive and wild populations in both the mycobiome and microbiome (Figure 2; PERMANOVA, *p* < 0.01). However, we found no significant differences in alpha diversity associated with captivity for either (Figure S4; Kruskal-Wallis, *p* > 0.05). When looking for ASVs that significantly differed with captivity using DESeq2, we found 21 fungal ASVs and one bacterial ASV that significantly decreased in abundance in captive individuals compared to their wild counterparts (Figure 3, Table 2). Fungal ASVs with the largest log_2_fold changes include members of the genera *Neodidymelliopsis, Aureobasidium, Neosetophoma, Dioszegia, Metschnikowia, Cladosporium, Buckleyzyma,* and an unclassified Cystobasidiomycetes species. For those ASVs with predicted fungal trophic modes, the majority were saprotrophic, with only a few ASVs having insect- or plant-pathotrophic predictions. The single bacterial ASV with a moderate decrease in abundance represents *Staphylococcus aureus*. We identified no ASVs in the mycobiome or microbiome with increased abundance in captive individuals.

**Figure 2.**
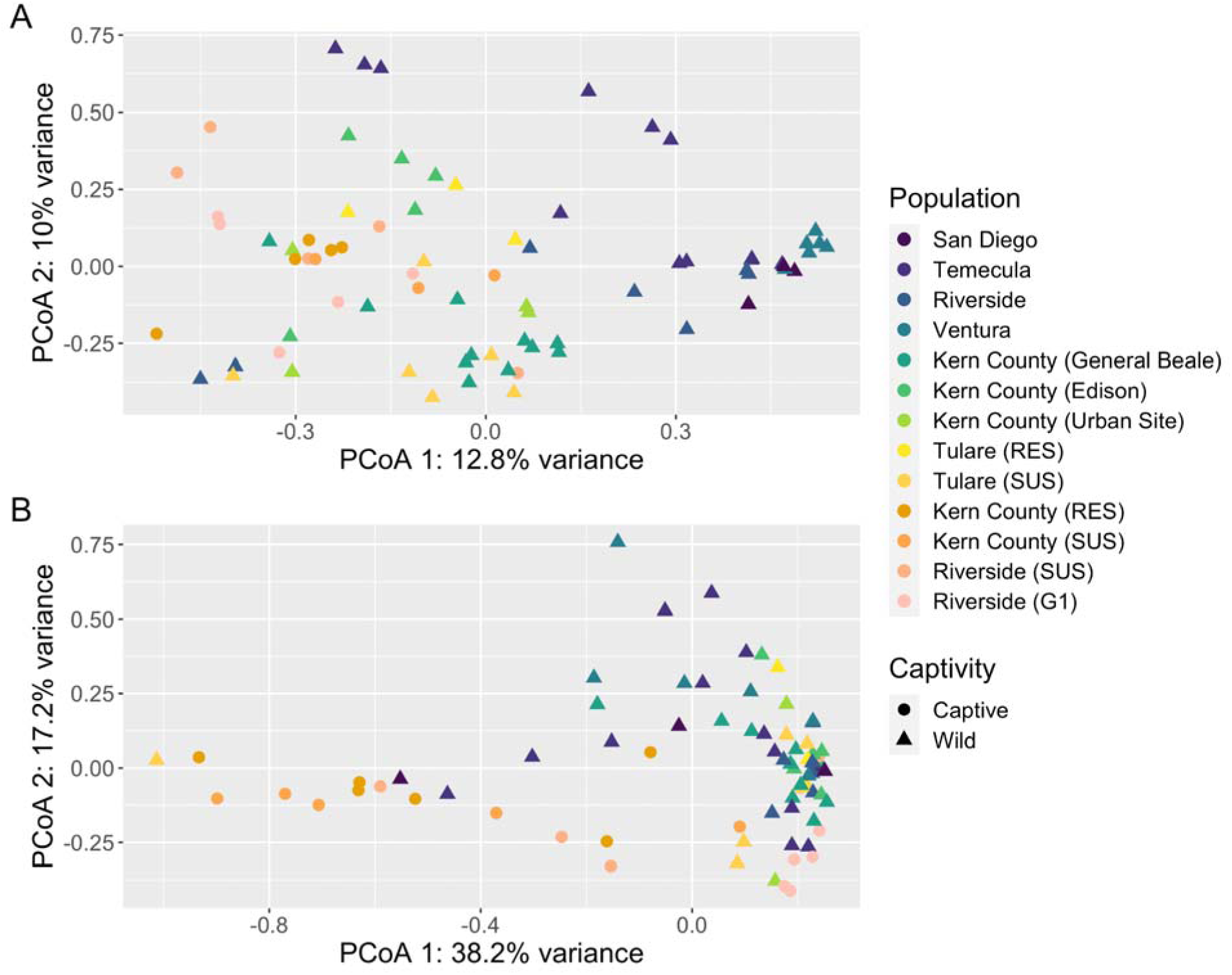
Community structure differs across geography and captivity. Principal-coordinate analysis (PCoA) visualization of Hellinger distances of (A) fungal and (B) bacterial communities. Individual GWSS are colored by population (collection site), with wild populations ordered by latitude and followed by captive populations, and have shapes based on captivity status.

**Figure 3.**
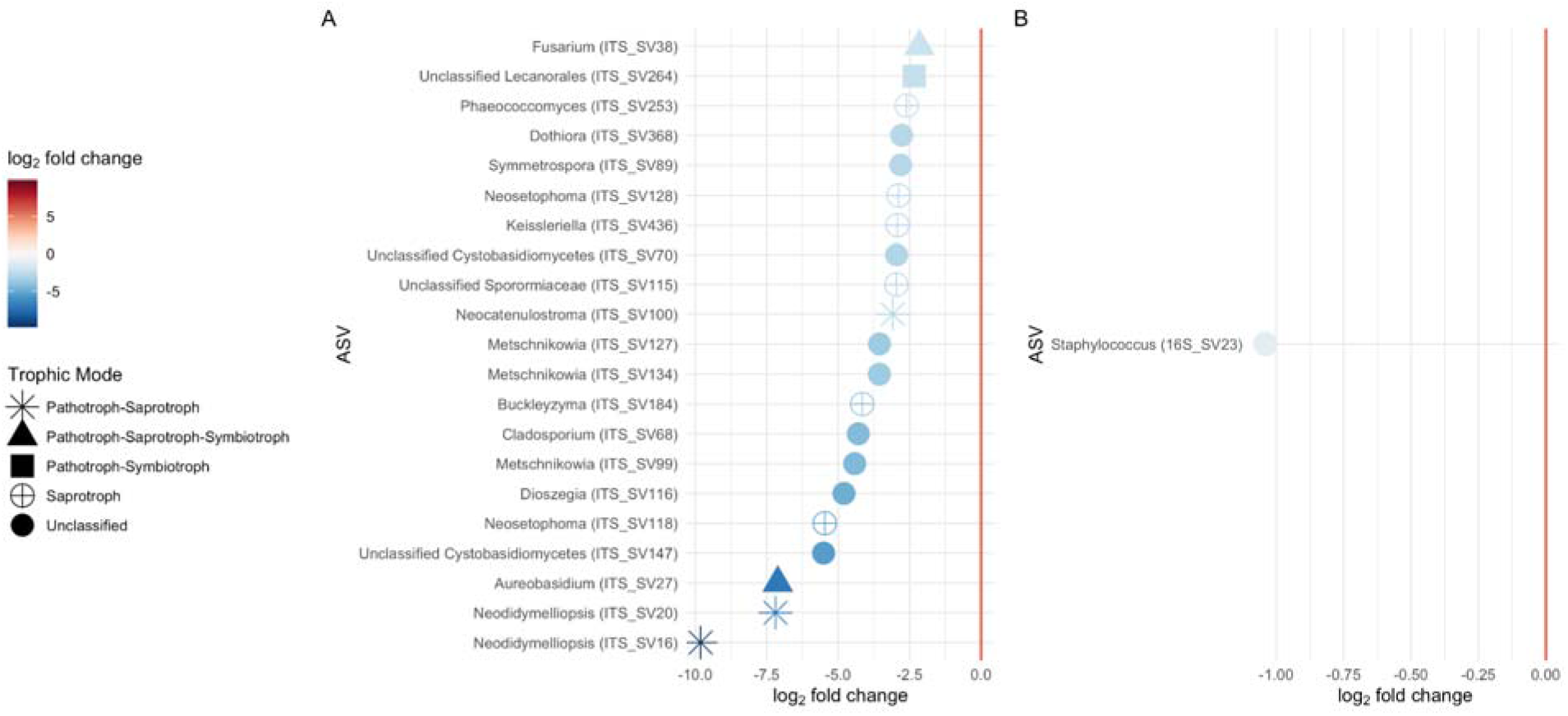
Differentially abundant ASVs associated with captivity status. Log_2_fold changes of significantly differentially abundant ASVs associated with captivity status, colored by log_2_fold change, and labeled by ASV and putative genera. For ITS1 ASVs, predicted trophic modes from FUNGuild are shown as shapes. A positive log_2_fold change indicates the ASV had higher abundance in captive individuals, while a negative log_2_fold change indicates it was higher in wild individuals. Full taxonomy for each ASV can be found in Table 2.

**Table 2.**
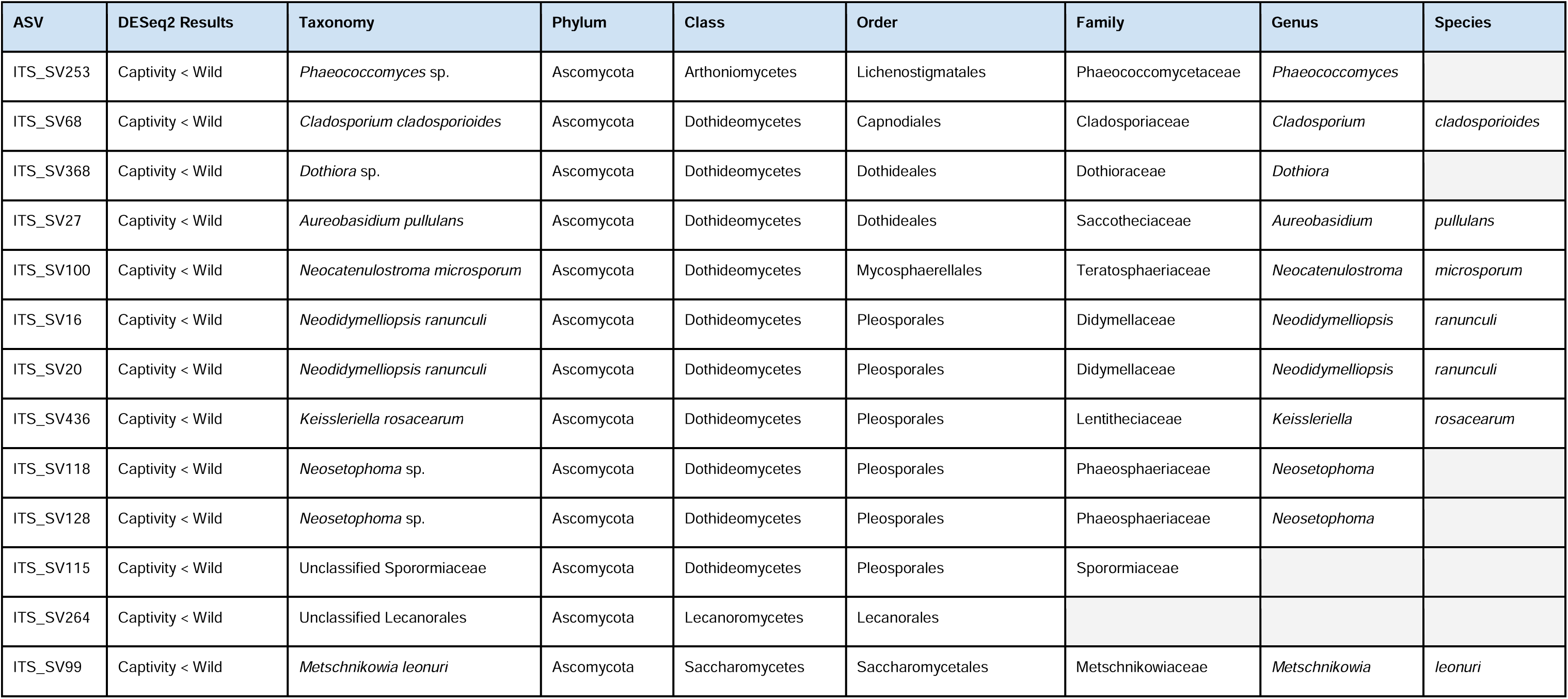
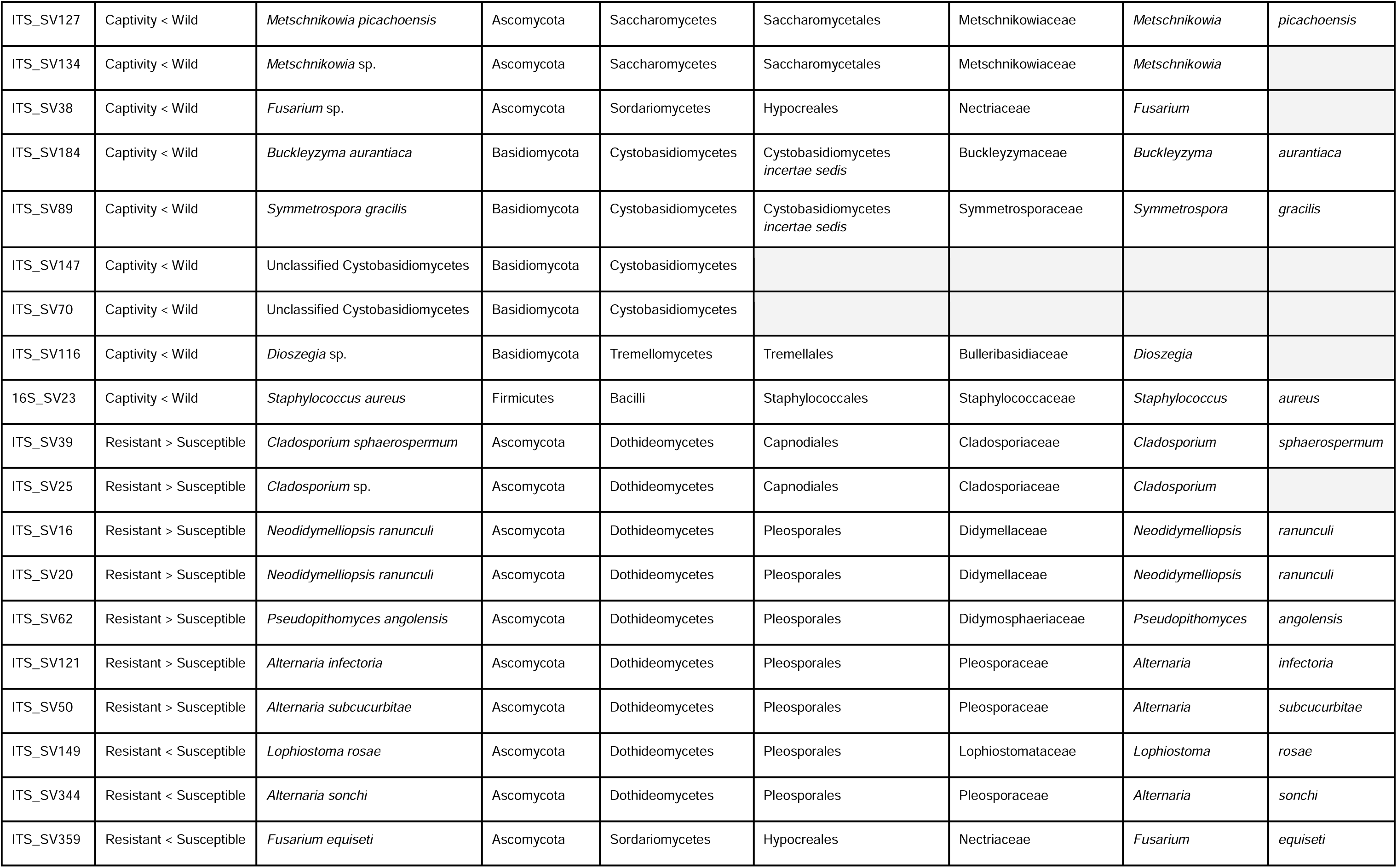
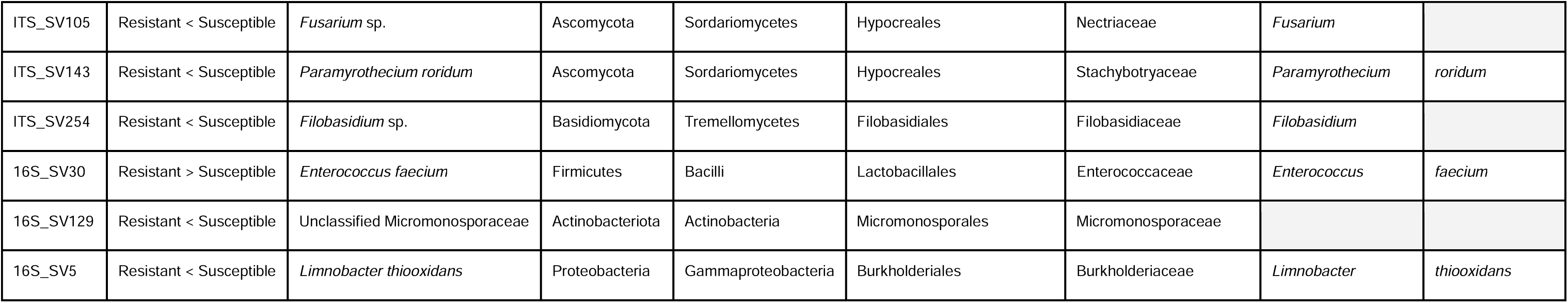
Differentially abundant ASVs. DESeq2 was used to identify ASVs that were differentially abundant between captive and wild-caught GWSS, and between insecticide-resistant wild-caught and insecticide-susceptible wild-caught GWSS. Here, for each predicted ASV that was differentially abundant between groups of interest, we report the ASV, the significant pairwise differential abundance comparisons (e.g., captivity < wild means that the ASV was in significantly higher abundance when associated with wild GWSS than with captive GWSS), and the predicted taxonomy of the ASV.

### Possible evidence of microbial signal of host insecticide-resistance in myco- or microbiomes

When testing for an association between community structure and host insecticide-resistance status, we found significant differences in beta diversity for the mycobiome (Figure S5; PERMANOVA, *p* = 0.02), but not the microbiome (*p* > 0.05). Host insecticide-resistance status explained a smaller proportion of mycobiome variation (*R^2^* = 0.05) compared to the variation explained by geographic region (*R^2^*= 0.15) or collection year (*R^2^* = 0.12). Further, for both the myco- and microbiomes, we found no differences in dispersion (betadisper, *p* > 0.05) or alpha diversity associated with host insecticide-resistance status (Figure S6; Kruskal-Wallis, *p* > 0.05).

While we only detected a possible association with host insecticide-resistance status in the overall community structure of the mycobiome, we still attempted to identify specific ASVs in both the microbiome and mycobiome that might correlate with host insecticide-resistance status using DESeq2. We identified significant log_2_fold change differences that were associated with host insecticide-resistance status in 13 ASVs in the mycobiome and three ASVs in the microbiome (Figure 4, Table 2). Seven fungal ASVs were higher in abundance on resistant individuals, with the largest log_2_fold changes observed from ASVs representing the genera *Alternaria* and *Pseudopithomyces*, while six fungal ASVs were higher in abundance on susceptible individuals with the largest log_2_fold changes in the genera *Lophiostoma*, *Filobasidium*, and *Paramyrothecium*. For ASVs with predicted fungal trophic modes, the majority were animal or plant pathotrophs, with some having saprotroph predictions. One bacterial ASV, representing the genus *Enterococcus*, was enriched in abundance on resistant individuals, while two bacterial ASVs, representing the genus *Limnobacter* and an unclassified member of the Micromonosporaceae family, were enriched on susceptible individuals.

**Figure 4.**
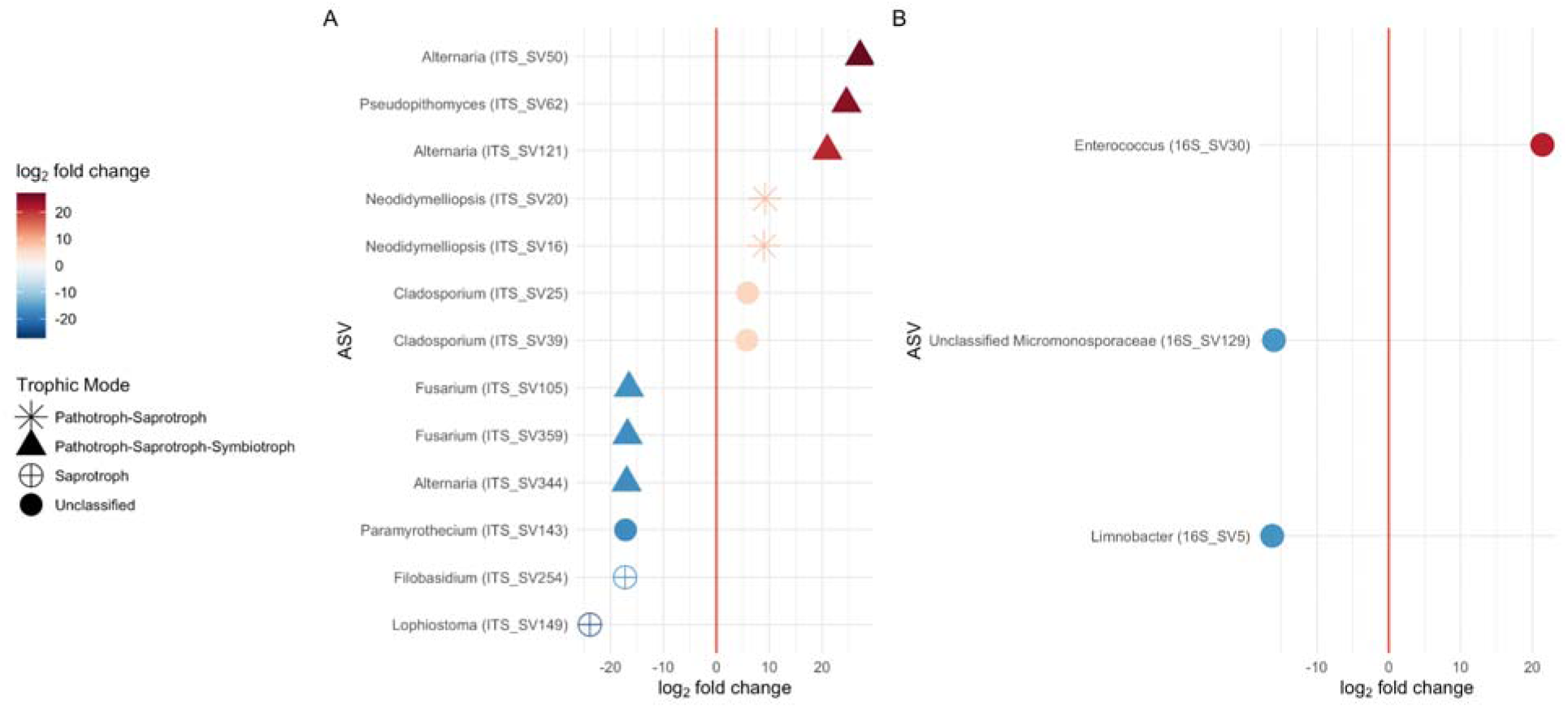
Differentially abundant ASVs associated with host insecticide-resistance status. Log_2_fold changes of significantly differentially abundant ASVs associated with host insecticide-resistance status of wild sharpshooters, colored by log_2_fold change, and labeled by ASV and putative genera. For ITS1 ASVs, predicted trophic modes from FUNGuild are shown as shapes. A positive log_2_fold change indicates the ASV had higher abundance in insecticide-resistant hosts, while a negative log_2_fold change indicates it was higher in insecticide-susceptible hosts. Full taxonomy for each ASV can be found in Table 2.

## Discussion

This study is the first to characterize the mycobiome of GWSS, and the first to profile both the microbiome and mycobiome across Southern California. We found that the mycobiome was dominated by putative plant pathogens and saprotrophs, while the microbiome was dominated by obligate and facultative symbionts. While the structure of the mycobiome varied across Southern California, the structure of the microbiome did not. Further, we observed a correlation between captivity and the structure of both the mycobiome and microbiome, and identified specific ASVs that were enriched in wild individuals versus their captive counterparts. Finally, we found that the structure of the mycobiome, but not the microbiome, was correlated with host insecticide-resistance status in wild GWSS, and we were able to identify members of both the mycobiome and microbiome that significantly varied in relative abundance with host insecticide-resistance status.

### The microbiome is dominated by obligate and facultative symbionts and is less diverse than the mycobiome

Obligate symbionts, *Ca. Baumannia cicadellinicola* and *Ca. Sulcia muelleri*, and the facultative symbiont *Wolbachia* sp. were the most abundant bacteria in the microbiome (Figure 1B). Given their importance to host metabolism, obligate symbionts have unsurprisingly been reported to dominate the bacterial community in other GWSS microbiome studies [37, 38]. While GWSS is known as a vector of *X. fastidiosa,* no ASVs representing *Xylella* were detected in the microbiome here, possibly due to our choice to sample whole insects. Our results are similar to other work from whole insects [11] but contrast with results from dissected foregut tissues [13]. Despite being only a facultative symbiont, *Wolbachia* sp. were detected in the majority (96.6%) of insects, which is similar to the high prevalence reported in other studies [13, 37]. Given the impact of *Wolbachia* sp. on reproductive fitness of insects in other species, its prevalence here in invasive GWSS in Southern California may be worth taking into account when considering GWSS biocontrol [76, 77].

While symbionts dominated the bacterial community, we also observed low abundances of rare genera including *Limnobacter*, *Acinetobacter*, *Methylophilus*, and *Pseudomonas,* many of which have been reported in association with GWSS previously [13, 36, 37]. It has been suggested that these rare taxa may be locally acquired through travel to and feeding on local plant hosts [37]. For example, *Pseudomonas* has been proposed as a core member of the grape endosphere [78]. However, we found no evidence of an association between geographic region and microbiome structure here similar to what has been found in studies of other invasive GWSS in California [13], but different from what has been reported for native GWSS in Texas [37]. We also found that the microbiome was less diverse than the mycobiome, which is consistent with descriptions from other insects including planthoppers and caterpillars [79, 80].

### Mycobiome varies across regions and may reflect local environment

In contrast with the microbiome, we found that the structure of the mycobiome was variable across geographic regions. A similar pattern has previously been reported for invasive beetles, where local habitat was found to strongly correlate with mycobiome structure [81], as well as for planthopper mycobiomes, which have been reported to vary across sampling sites [80]. The taxonomic composition of the mycobiome appears dominated by putative plant- and soil-associated pathogens and saprotrophs, similar to findings from planthoppers where it has been suggested that the mycobiome is acquired from the environment [82]. In support of this, a majority of the fungal genera found associated with GWSS here have also been previously reported in association with plants and rhizosphere soil in vineyards and orchards [78, 83–89], and the vineyard mycobiome has been found to also vary with geography [90]. Thus, the prevalence of putative plant associated fungi in the GWSS mycobiome suggests a possible role for diet as a critical origin source for the fungal community of insects [82]; however, a study comparing the caterpillar-gut mycobiome to leaf communities found them to be distinctly structured, as compared to the caterpillar microbiome which has been reported to mirror its diet, indicating that diet alone may not fully explain mycobiome acquisition in pest insects [79, 91].

### Captivity may lead to differences in both the myco- and microbiomes

Many studies have reported microbial community differences related to captivity in mammals [92, 93], birds [94, 95], amphibians [96, 97], and also in insects including beetles [98], armyworms [99], and fruit flies [100]. Often these studies report a reduction of alpha diversity [101, 102], which we did not observe here; instead we found a reduction of specific ASVs associated with captivity (Figure 3). Captivity involves changes in many factors which likely impact microbial communities, including dietary restrictions, habitat changes (e.g., enclosed space, stable temperature, constant light), reduced species interactions, and exposure to human-associated microbes. Associations between the microbiome and captivity have been suggested to be due to the changes in diet, behavior and environment compared to wild individuals [101–103]. Given our findings on the possible link between the mycobiome and local habitat, it is possible the fungal taxa observed to be less abundant here are simply missing from the captive diet or habitat, which is limited in comparison to their wild counterparts. Most of the fungal genera identified as having lower abundance in the captive populations have been reported as putative plant saprotrophs [104, 105], insect pathogens [106], plant surface-associated [107, 108] or phytopathogens, including of roses [105, 109], citrus [110], and other flowering plants [111]. Therefore, it is certainly possible that captive GWSS are not being exposed to these fungal sources, and thus that is the reason these specific ASVs are lower in abundance in captive GWSS. Future work could test the effect of altered diet on fungal community assembly in GWSS through dietary manipulations in captivity to assess the utility of GWSS as a possible indicator species of plant fungal disease in wild populations.

In the microbiome of captive individuals, we saw only a significant, but relatively small, reduction in *Staphylococcus*, which has been previously part of the natural microbiome of the sharpshooter, *Acrogonia citrina* [112]. In contrast to our findings, *Staphylococcus* abundance was found to be higher in abundance in captive beetles compared to their wild counterparts [98]. While it is unclear why *Staphylococcus* is lower in abundance in captive GWSS, it is possible that similar to the mycobiome, this change relates to a captive diet or environment. Given the changes in the myco- and microbiomes between captive and wild populations, future work should consider these microbial communities in any future GWSS biocontrol strategies and investigate possible benefits of ‘rewilding’ these communities with environmentally-acquired microbes prior to release [113, 114].

### Members of the myco- and microbiomes are correlated with host insecticide-resistance status

Here we found evidence of a link between mycobiome structure and host insecticide-resistance status and were able to identify specific ASVs in both the bacterial and fungal communities that correlated with host insecticide-resistance status. Similar correlations between host insecticide-resistance and the microbiome has been previously reported in the brown planthopper [115], the cotton bollworm [116, 117], mosquitoes [118], and cockroaches [119]. Many of the fungal ASVs that correlated here with host insecticide-resistance status are predicted to be plant-associated or phytopathogens [89, 109, 120, 121], again indicating the possible importance of environmental acquisition through shared space or diet. Given that historical use of insecticides against GWSS in Southern California has involved area-wide treatments [17, 18], the plant-associated microbes in these regions are also being subjected to these insecticides; thus, it is possible that these microbes are more abundant on leaves in these regions given their own ability to resist or detoxify these compounds, and GWSS may just be acquiring them from the local environment.

While overall microbiome structure was not correlated with host insecticide-resistance status, we did identify a few bacterial ASVs with differential abundance between resistant and susceptible GWSS. Of these, most have been previously reported in insect guts [122–124]. However, only *Enterococcus* was found to be in higher abundance in insecticide-resistant GWSS microbiomes. In diamondback moths, *Enterococcus* sp. were found to enhance insecticide-resistance, which the authors hypothesized was due to gut bacteria preventing or restoring damage done to the host immune system by insecticides [27]. It is possible that *Enterococcus* has a similar beneficial role here for GWSS. Future work should isolate the bacteria and fungi from GWSS and host plants from insecticide-resistant locations and assess whether these microbes confer resistance *in vitro* and then whether they confer resistance after inoculation into sharpshooters *in vivo*.

## Conclusion

Overall this study surveys the microbiome and mycobiome associated with invasive GWSS from across Southern California, serving as the first characterization of the fungal community associated with GWSS. We identified significant differences in community structure between locations for the mycobiome, but not the microbiome, indicating environmental acquisition, possibly through diet, of the GWSS fungal community. We also found that captivity is correlated with changes in the structure of the fungal and bacterial communities associated with GWSS, with some members of the communities found to be less abundant in captive populations. We found a potential association with host insecticide-resistance status and mycobiome structure in wild GWSS and were able to identify specific ASVs correlated with host insecticide-resistance status in both the microbiome and mycobiome. This study provides foundational insight into the mycobiome and microbiome of GWSS across Southern California and serves as additional support for a growing body of literature surrounding the effects of captivity on host-associated microbial communities. The differences in the microbial communities between field and laboratory-reared GWSS will be an important consideration in genetic control-based strategies in which gene-edited, laboratory-reared GWSS should not be at a competitive disadvantage to the target field population. Finally, this work identifies a possible link between members of both the bacterial and fungal communities and host insecticide-resistance status which should be explored further.

## Supporting information

Suppmental Materials including Tables S1, S2 and Figures S1-S6

## Data availability

Sequence reads generated for the ITS1 region and the 16S rRNA gene libraries were deposited in GenBank under BioProject ID <u>PRJNA934966</u>. Analysis scripts are available on Github (https://github.com/stajichlab/GWSS_Microbiome_Analysis) and archived in Zenodo [125].

## Acknowledgments

We would like to thank members of the Byrne lab, specifically Tim Roose and Johnny Sun, for their contributions to the field collections of GWSS used in this work.

## Author contributions

C.L.E. and J.E.S. conceived the idea. L.L.W., P.W.A., R.R. and J.E.S. obtained funding. F.J.B. and D.J.B. provided samples. D.J.B. and B.G.V. maintained captive lines. C.L.E., J.W.W., T.K., and I.D.S.P. performed molecular analysis. C.L.E., J.W.W. and T.K. prepared sequencing libraries. C.L.E. analyzed the data and wrote the manuscript. All authors read, reviewed, and edited the manuscript.

