## Supplementary material for "Geographical Survey of the Mycobiome and Microbiome of Southern California Glassy-winged Sharpshooters": Suppmental Materials including Tables S1, S2 and Figures S1-S6

**Supplemental Tables and Figures**

**Table S1.** Pairwise PERMANOVA contrast tests across geographic regions on the mycobiome. Here we report the corrected p-values from pairwise PERMANOVA contrast tests which were used to assess which pair-wise regional comparisons were driving observed differences in mean centroids. Pairwise comparisons that are significantly different are bolded (*p* < 0.05) and shaded gray.

|  | Temecula | Riverside | Ventura | Kern County | Tulare |
| --- | --- | --- | --- | --- | --- |
| San Diego | **0.025** | 0.079 | **0.021** | **0.003** | **0.008** |
| Temecula | NA | **0.011** | **< 0.001** | **< 0.001** | **< 0.001** |
| Riverside | NA | NA | **0.002** | **0.023** | **0.048** |
| Ventura | NA | NA | NA | **< 0.001** | **0.001** |
| Kern County | NA | NA | NA | NA | 0.381 |

**Table S2.** Tukey *Post-hoc* tests of dispersion across geographic regions on the mycobiome. Here we report the corrected p-values from Tukey *Post-hoc* tests which were used to assess which pair-wise regional comparisons were driving observed dispersion differences. Pair-wise comparisons that are significantly different are bolded (*p* < 0.05) and shaded gray.

|  | Temecula | Riverside | Ventura | Kern County | Tulare |
| --- | --- | --- | --- | --- | --- |
| San Diego | **0.025** | **0.005** | 0.999 | **< 0.001** | **0.002** |
| Temecula | NA | 0.857 | **0.007** | 0.105 | 0.692 |
| Riverside | NA | NA | **0.001** | 0.931 | 1.000 |
| Ventura | NA | NA | NA | **< 0.001** | **< 0.001** |
| Kern County | NA | NA | NA | NA | 0.965 |

**Figure S1.** Map depicting collection regions. Map diagram showing California with labels depicting different collection regions.
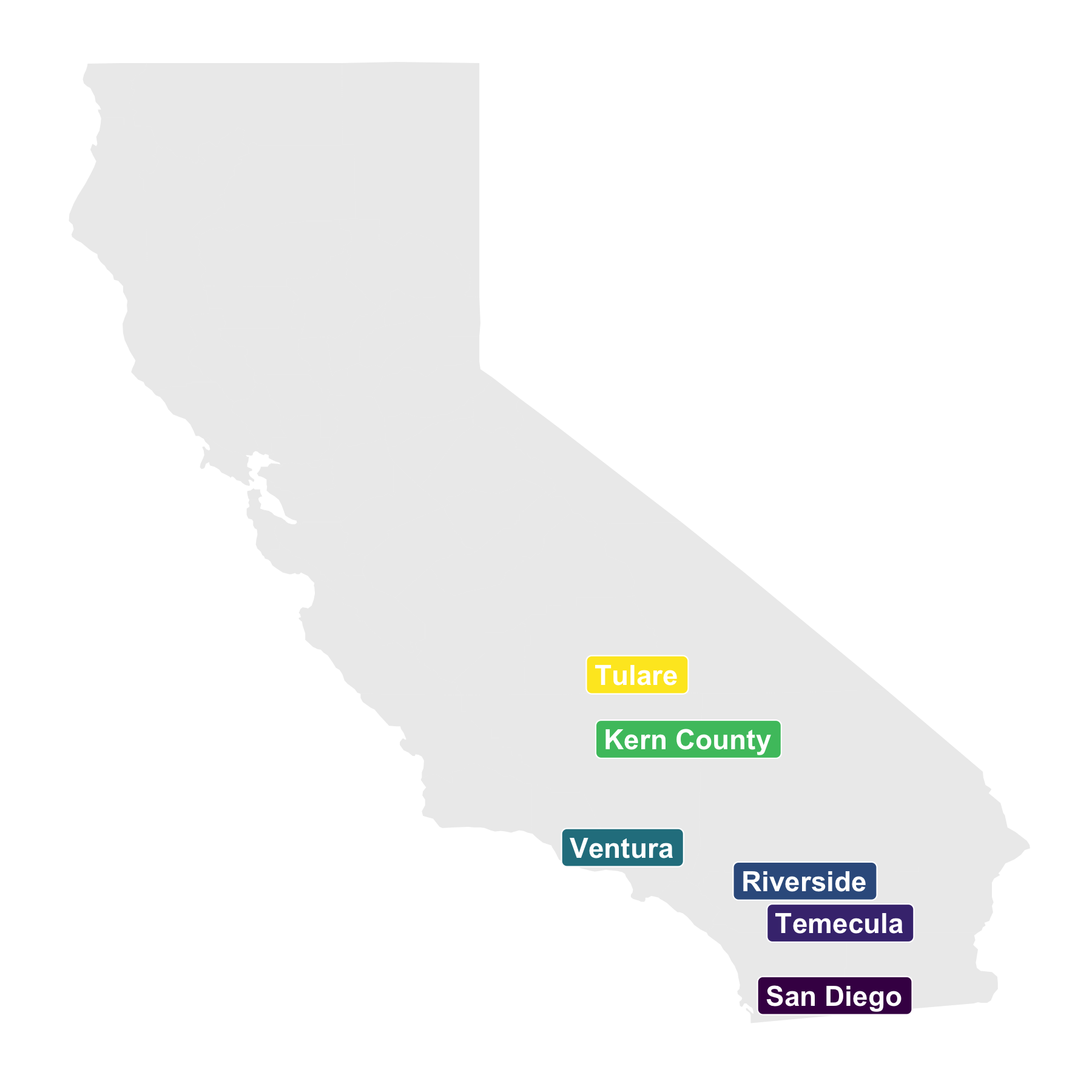

**Figure S2.** Community structure of wild sharpshooters across space and time. Principal-coordinate analysis (PCoA) visualization of Hellinger distances of (A) fungal and (B) bacterial communities, with colors indicating collection region and shapes indicating collection year.

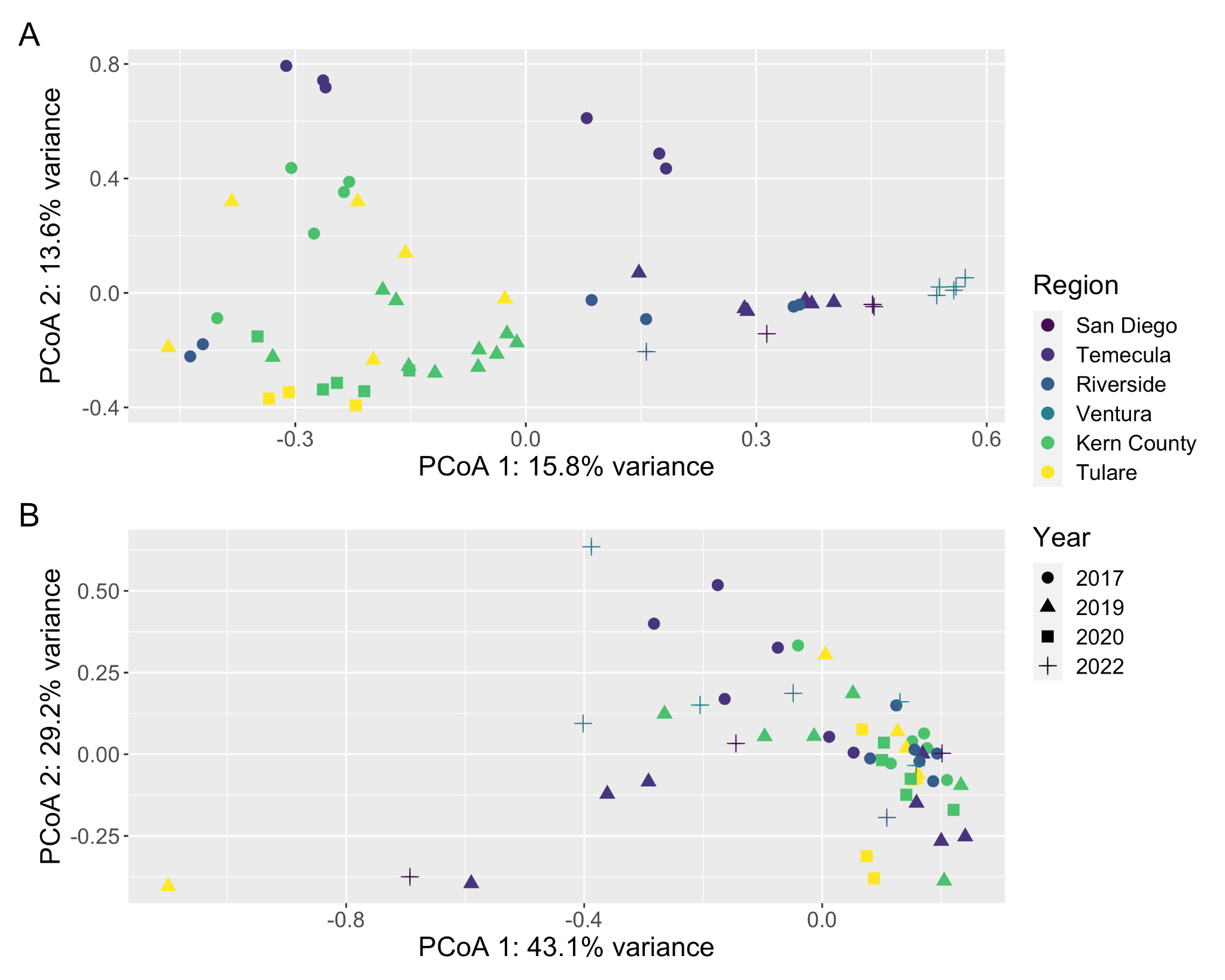

**Figure S3.** Alpha diversity does not vary across wild populations. Here we depict barcharts of Shannon diversities for the (A) ITS1 region and (B) 16S rRNA gene datasets. The standard error of the mean Shannon diversity of each population is represented by an error bar, and bars are colored by population.

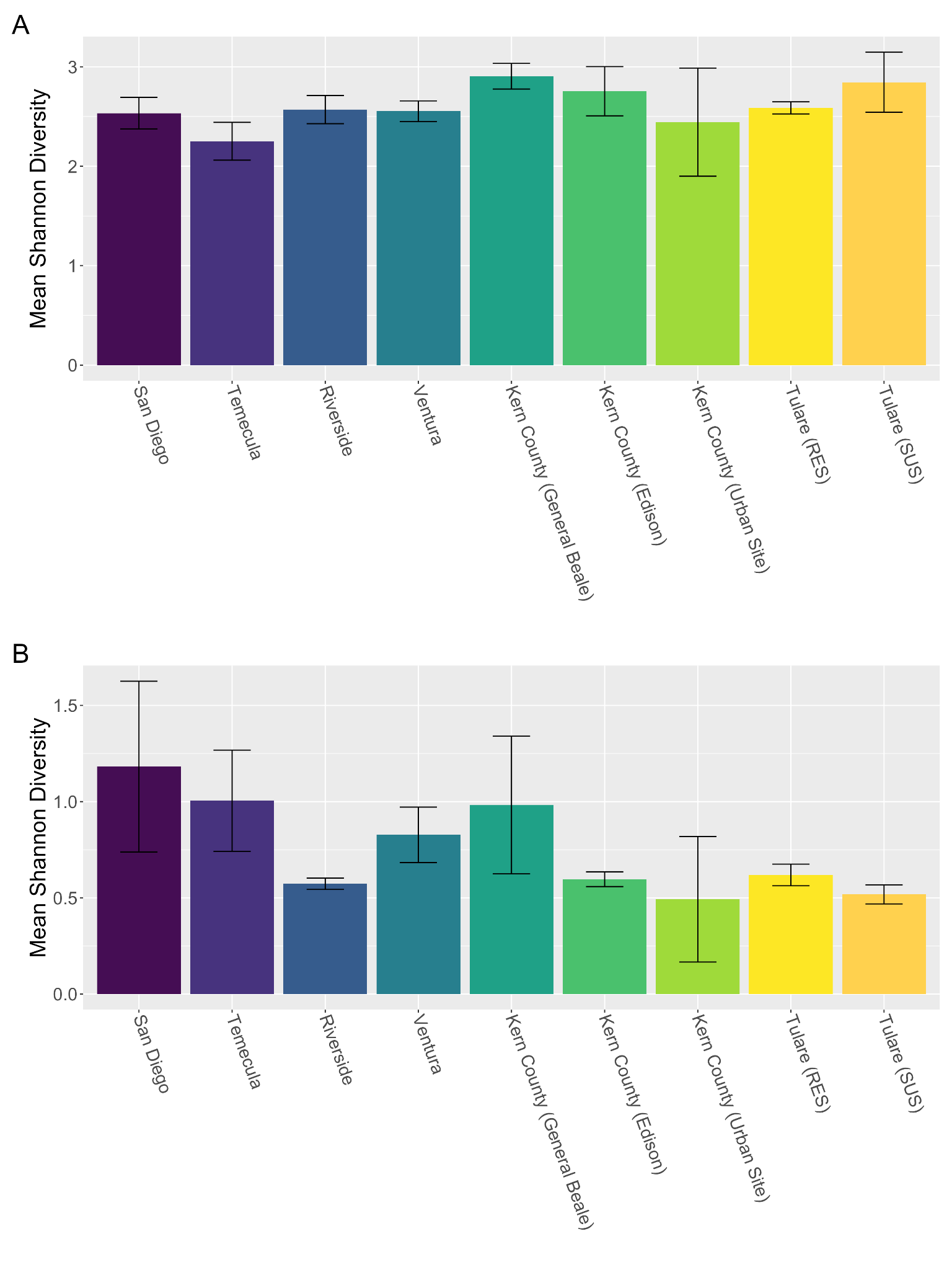

**Figure S4.** Alpha diversity does not vary with captivity. Here we depict barcharts of Shannon diversities for the (A) ITS1 region and (B) 16S rRNA gene datasets. The standard error of the mean Shannon diversity for captive and wild sharpshooters is represented by an error bar, and bars are colored by captivity status. Comparisons that are significantly different from each other are notated by different letters (e.g., a vs. b).

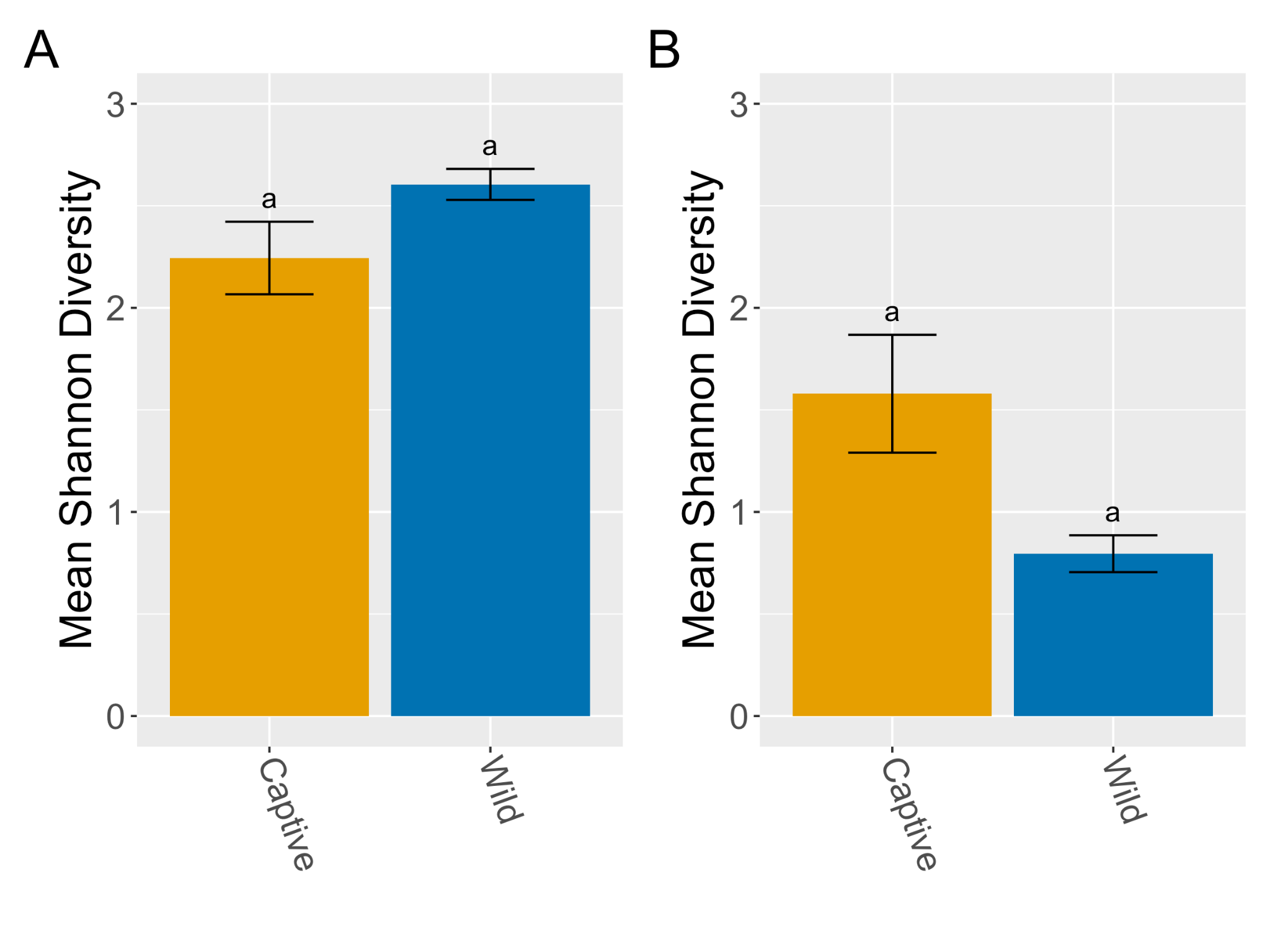

**Figure S5.** Community structure of wild sharpshooters with varying host insecticide-resistance status. Principal-coordinate analysis (PCoA) visualization of Hellinger distances of (A) fungal and (B) bacterial communities, with colors and shapes indicating host insecticide-resistance status.

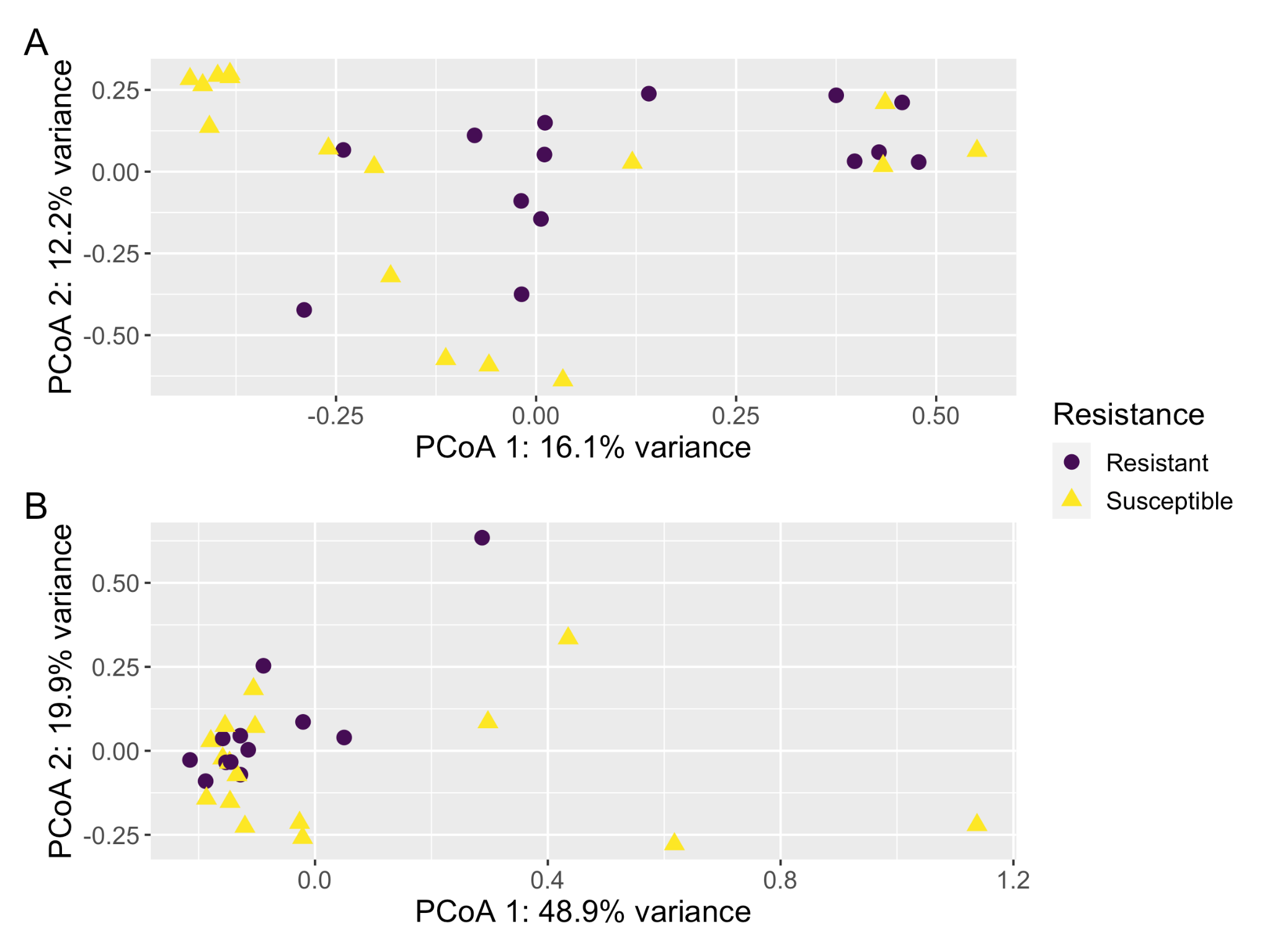

**Figure S6.** Alpha diversity does not vary with host insecticide-resistance status. Here we depict barcharts of Shannon diversities for the (A) ITS1 region and (B) 16S rRNA gene datasets. The standard error of the mean Shannon diversity for insecticide-resistant and susceptible hosts is represented by an error bar, and bars are colored by host insecticide-resistance status. Comparisons that are significantly different from each other are notated by different letters (e.g., a vs. b).

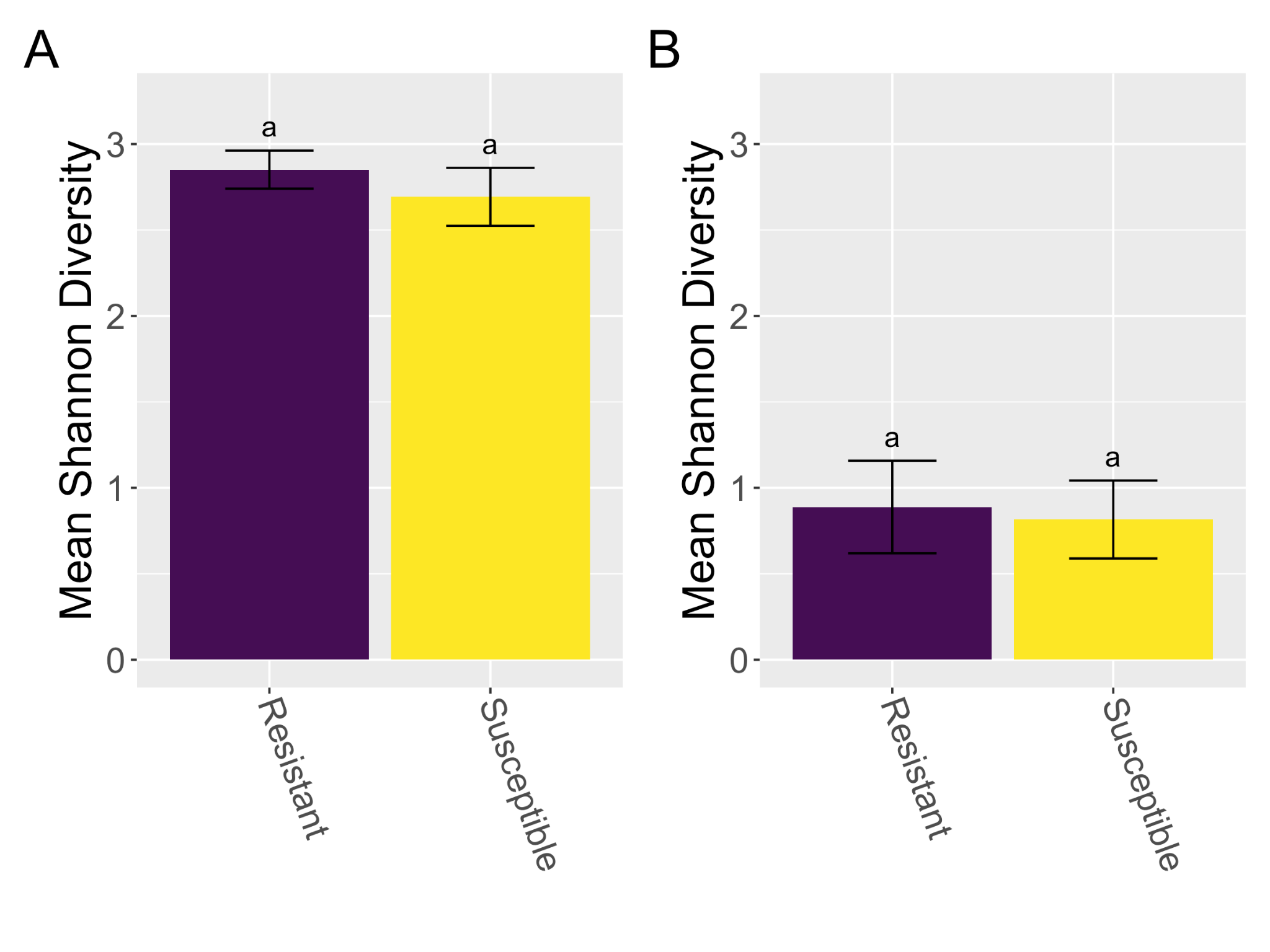
